# Phylogenomics reveals an almost perfect polytomy among the almost ungulates (*Paenungulata*)

**DOI:** 10.1101/2023.12.07.570590

**Authors:** Jacob Bowman, David Enard, Vincent J. Lynch

## Abstract

Phylogenetic studies have resolved most relationships among *Eutherian* Orders. However, the branching order of elephants (*Proboscidea*), hyraxes (*Hyracoidea*), and sea cows (*Sirenia*) (i.e., the *Paenungulata*) has remained uncertain since at least 1758, when Linnaeus grouped elephants and manatees into a single Order (*Bruta*) to the exclusion of hyraxes. Subsequent morphological, molecular, and large-scale phylogenomic datasets have reached conflicting conclusions on the branching order within *Paenungulates*. We use a phylogenomic dataset of alignments from 13,388 protein-coding genes across 261 *Eutherian* mammals to infer phylogenetic relationships within *Paenungulates*. We find that gene trees almost equally support the three alternative resolutions of *Paenungulate* relationships and that despite strong support for a *Proboscidea*+*Hyracoidea* split in the multispecies coalescent (MSC) tree, there is significant evidence for gene tree uncertainty, incomplete lineage sorting, and introgression among *Proboscidea*, *Hyracoidea*, and *Sirenia*. Indeed, only 8-10% of genes have statistically significant phylogenetic signal to reject the hypothesis of a *Paenungulate* polytomy. These data indicate little support for any resolution for the branching order *Proboscidea*, *Hyracoidea*, and *Sirenia* within *Paenungulata* and suggest that *Paenungulata* may be as close to a real, or at least unresolvable, polytomy as possible.

## Introduction

Systematic studies have successfully resolved most of the phylogenetic relationships within *Eutherian*^1^ mammals. Some clades, however, have remained recalcitrant to phylogenetic resolution, including the interrelationships between elephants (*Proboscidea*), hyraxes (*Hyracoidea*), and manatees (*Sirenia*). While these lineages were combined into the superorder *Paenungulata*, the “almost ungulates”, by Simpson (1945), who also noted the possibility that *Hyracoidea* might be more closely related to the ungulate order *Perissodactyla*, the recognition that *Proboscideans* and *Sirenians* are closely related to each other is much older (**Figure 1A**). In *Systema Naturae* (1758), for example, Linnaeus classified elephants and sirenians together under the name *Bruta* (Linné, 1758); this group also included sloths, anteaters, and pangolins, anticipating the close phylogenetic relationships of *Paenungulates* and *Xenarthrans*. Similarly, de Blainville (1836) classified elephants and manatees together in the group “*les gravigrades*”, but he did not formalize this proposal (Blainville, 1839).

**Figure 1.**
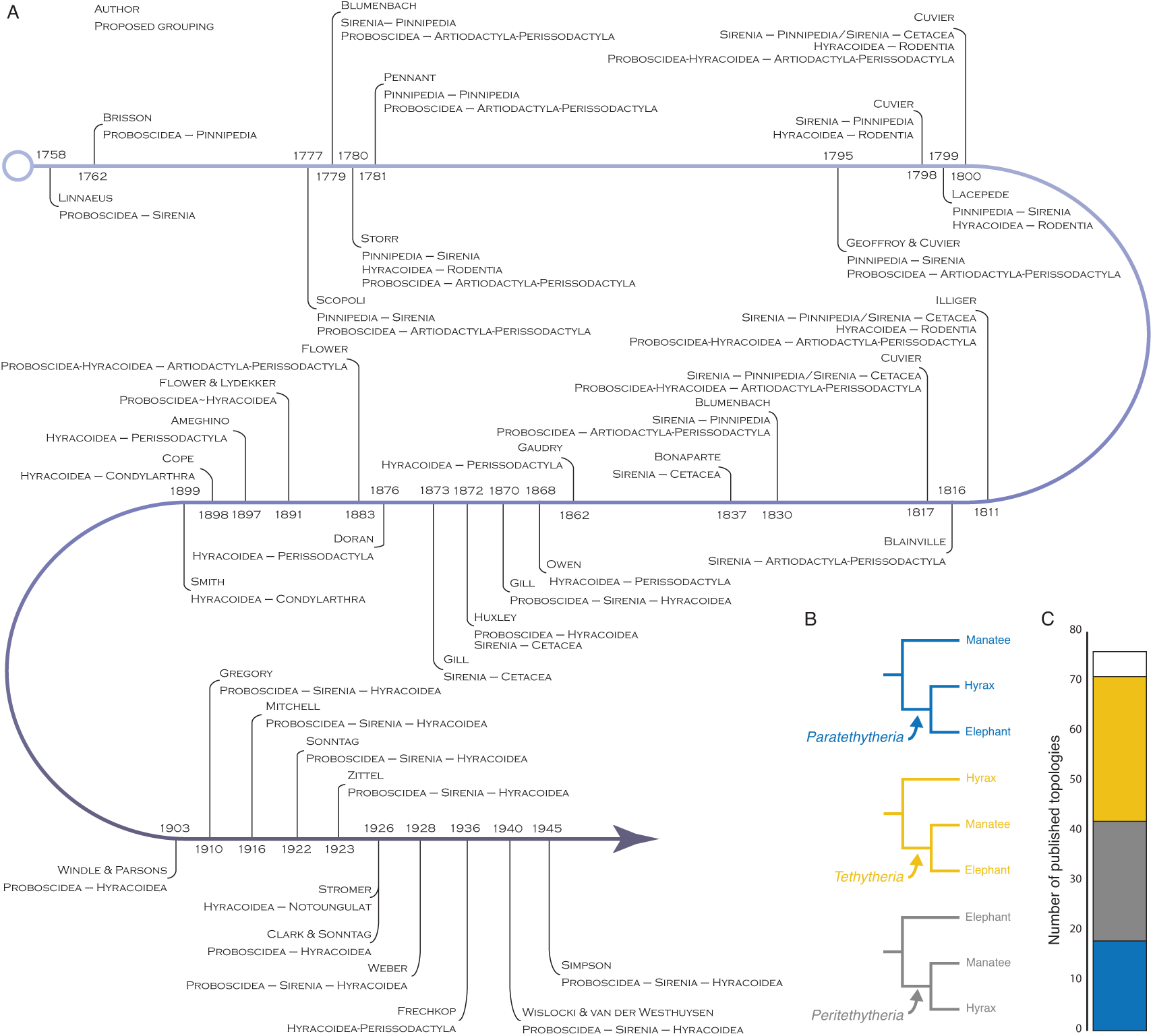
Historical and recent hypothesis of phylogenetic relationships among Proboscidea, Hyracoidea, Sirenia. **A.** Timeline (Linnaeus to Simpson) showing proposed relationships between *Proboscidea*, *Hyracoidea*, *Sirenia,* and other mammalian Orders. Modified from (J. H. Shoshani, 1986) Table 1. **B.** The three alternative resolutions of *Paenungulata.* We refer to the *Proboscidea*+*Hyracoidea* clade as *Paratethytheria*, the *Sirenia*+*Hyracoidea* clade as *Peritethytheria*, and the *Proboscidea*+*Sirenia* clade as *Tethytheria*. **C.** Stacked bar chart showing the number of published molecular phylogenies supporting the three alternative resolutions of *Paenungulata* shown in panel B. Bar colors follow the coloring in panel B. White represents published studies supporting an unresolved relationship.

**Figure 2.**
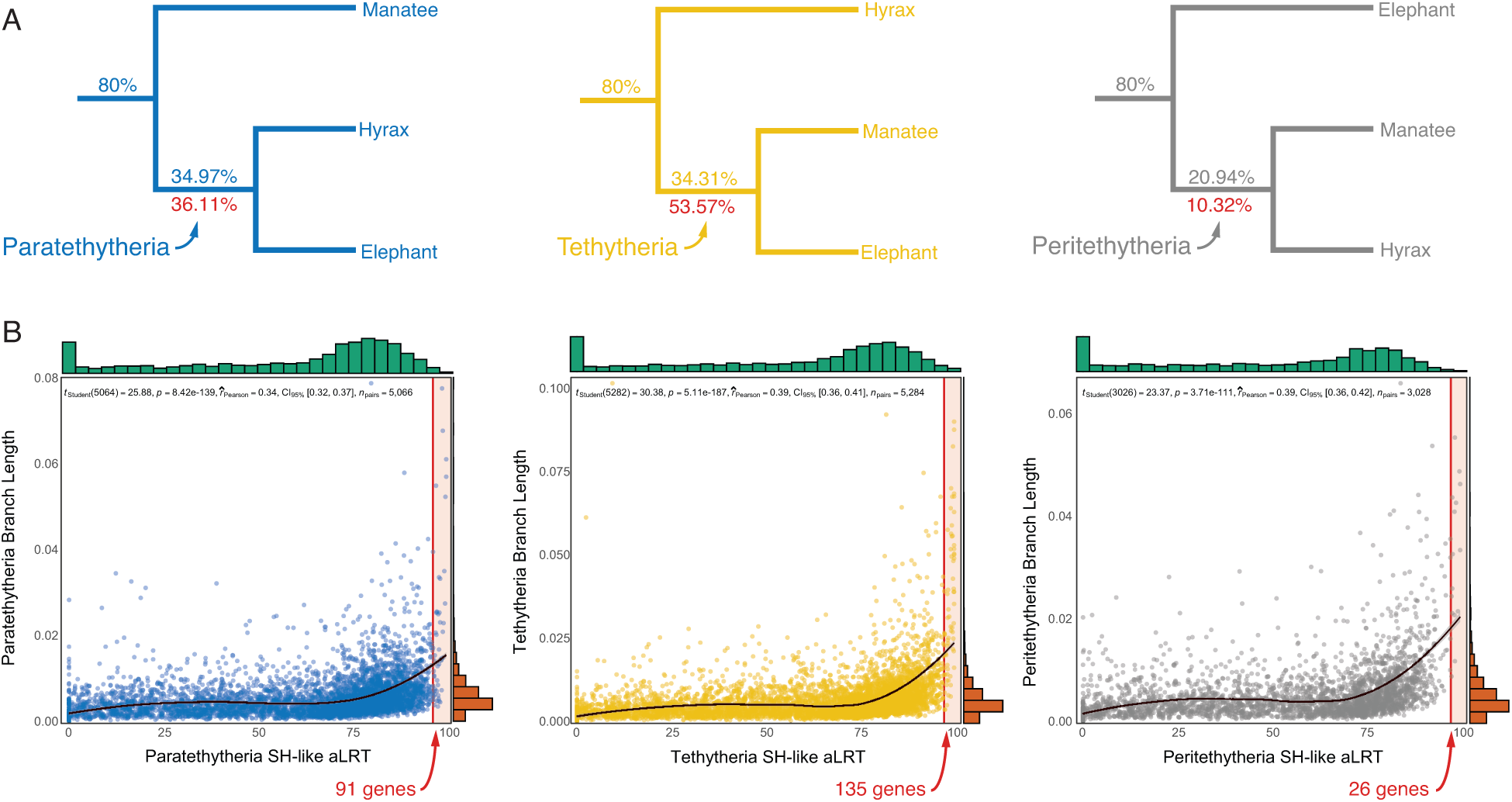
Gene trees are almost evenly divided between the three alternative resolutions of *Paenungulata*. **A.** The three alternative resolutions of *Paenungulata*, the percent of genes supporting each resolution are shown above each branch. The percent of genes with SH-like aLRT values ≥ 95% supporting each split is shown below branches in red. **B.** Scatterplot showing the correlation between branch length (substitutions per site) and SH-like aLRT support values for each gene supporting *Paratethytheria*, *Tethytheria*, and *Peritethytheria*. Loess regression is a black line (95% confidence interval around repression is shown in gray). The distribution of branch lengths and aLRT values are shown along the top (teal) and side (orange) of the plot, respectively. Vertical red lines indicate SH-like aLRT ≥ 95%, and the number of genes supporting *Paratethytheria*, *Tethytheria*, and *Peritethytheria* among genes with SH-like aLRT ≥ 95% is shown in red.

The clade uniting *Proboscideans* and *Sirenians* to the exclusion of other taxa was officially proposed and named *Tethytheria* by McKenna (1975), who, like Simpson also supported a close relationship between *Hyracoidea* and *Perissodactyla* (Domning et al., 1986; McKenna, 1975); Although McKenna (1975) does not explicitly describe the etymology of the name *Tethytheria*, it is likely named after the Tethys sea around which these lineages originated (Domning et al., 1986; McKenna, 1975; Prothero and Williams, 2017). Subsequently, Novacek (1982) proposed that *Hyracoidea* was a sibling lineage to the *Tethytheria* within *Paenungulata* (Novacek, 1982). The story of *Paenungulate* systematics, however, is much more complicated than this straightforward narrative suggests (J. H. Shoshani, 1986). Indeed, *Proboscidea*, *Hyracoidea*, and *Sirenia* have been considered closely related to numerous other mammalian orders, most often *Tethytheria* to *Pinnepedia* or *Cetacea*, and *Hyracoidea* to *Perissodactyla*. *Proboscidea*, *Hyracoidea*, and *Sirenia* were even considered to have close affinities to each other by several 19^th^ and early 20th-century morphologists (**Figure 1A**).

Molecular phylogenetic studies have almost uniformly agreed that *Proboscidea*, *Hyracoidea*, and *Sirenia* formed a monophyletic group within *Eutheria*. Still, they disagree on how these lineages are related to each other^2^. Indeed, while there are only three possible resolutions to how *Proboscidea*, *Hyracoidea*, and *Sirenia* can be related to each other (**Figure 1B**), each has received substantial support (**Figure 1C** and **Supplementary Table 1**). Even studies based on large multigene, phylogenomic, and rare genomic events such as chromosome rearrangements, transposable element insertions, gene duplications, and gene losses, infer conflicting relationships within *Paenungulates* (**Supplementary Table 1**). Therefore, we assembled a large dataset of alignments from 13,388 protein-coding genes across 261 *Eutherian* mammals, including an African elephant, rock and tree hyrax, and manatee, and used several complementary methods to infer their phylogenetic relationships. We found that gene trees were almost evenly divided among the three alternative *Paenungulate* resolutions and that relatively few genes had enough statistical support to favor one resolution over another. While the species tree favors the *Paratethytheria* tree, there is significant phylogenetic discordance among genes and sites resulting from incomplete lineage sorting, introgression, and gene tree uncertainty. Furthermore, tree topology tests cannot reject a *Paenungulate* polytomy for most genes (>90%). Thus, there is little support for any resolution for the branching order of *Proboscidea*, *Hyracoidea*, and *Sirenia,* and *Paenungulata* is as close to a real polytomy as possible.

## Methods

### Assembling alignments of orthologous protein-coding genes from 261 Eutherians

The alignment pipeline starts with Ensembl v99 human coding sequences, using the longest isoforms for each gene. We then search for the orthologs of these human coding sequences within the genome assemblies of 260 other mammals (**Supplementary Table 2**) with a contig size greater than 30kb in the NCBI assembly database as of July 2020. We select a minimum 30kb contig size to avoid excessive truncated orthologous coding sequences. To extract the orthologous coding sequences (CDS) to the human CDS, we use the best Blat reciprocal hits from the human CDS to each other mammalian genome and back to the human genome. We use Blat matching all possible reading frames, with a minimum identity set at 30% and the “fine” option activated (Kent, 2002). We excluded genes with less than 251 best reciprocal hits out of the 261 (human+other mammals) species included in the analysis. We find 13,491 human CDS genes with the best reciprocal hits orthologs in at least 250 other mammals; for reference, this corresponds to 68% of all protein-coding genes in the human genome.

We aligned each orthologous gene with Macse v2 (Ranwez et al., 2018). Macse v2 is a codon-aware multiple sequence aligner that can explicitly identify frameshifts and readjust reading frames accordingly. This feature of Macse is particularly important because erroneous indels introduced in coding sequences during genome sequencing and assembly processes can be common and cause frameshifts that many aligners do not consider. These sequencing and alignment errors can result in substantial misalignments of coding sequences due to incomplete codons. We start with the human CDS because they are likely the highest quality regarding sequencing and annotations. We use Macse v2 with maximum accuracy settings.

The alignments generated by Macse v2 were edited by HMMcleaner with default parameters (Franco et al., 2019). HMMcleaner is designed to remove, in a species-specific fashion, “fake” substitutions that are likely genome sequencing errors. HMMcleaner also removes “false exons” that might have been introduced during the Blat search. “Fake exons” are intronic segments that, by chance, are similar to exons missing from an assembly due to a sequencing gap. When looking for the most similar non-human CDS using Blat, Blat can sometimes “replace” the missing exon with a similar intronic segment.

After using HMMcleaner to remove suspicious parts of the CDS alignments in each species, we only select those still complete codons to remain in the alignments. In alignments of coding sequences, the flanks of indels usually include a higher number of misaligned substitutions. In each separate species, we further remove the x upstream or downstream codons from the alignments if more than x/2 of these codons code for amino acids that are different from the consensus amino acid in the whole alignment, with x varying from 1 to 20. For example, if eight of the 12 amino acids on the left side of an indel are different from the consensus in a given species, we remove the 12 corresponding codons. If two of the three amino acids on the right side of an indel are different from the consensus in a given species, we remove the three corresponding codons.

### Phylogenomic Analyses

We use IQTREE2 v.2.2.2 COVID-edition (Nguyen et al., 2015) to infer maximum likelihood phylogenetic gene trees from all 13,491 genes (nucleotide sequences) after the best-fitting model was identified for each gene by ModelFinder v.1.42 (Kalyaanamoorthy et al., 2017) with the –m MFP option; note that tree searches failed for 103 genes, thus our final dataset of trees includes 13,388 gene trees. Branch supports were assessed with the Shimodaira-Hasegawa-like approximate likelihood ratio (SH-like aLRT) test with 1000 replicates (Anisimova et al., 2006; Guindon et al., 2010) with the –alrt 1000 option. We also used IQTREE2 to perform tree topology tests with the RELL approximation (Kishino et al., 1990), including the bootstrap proportion, the Kishino-Hasegawa (Kishino and Hasegawa, 1989) and Shimodaira-Hasegawa tests (Shimodaira and Hasegawa, 1999), the weighted Kishino-Hasegawa and Shimodaira-Hasegawa tests, expected likelihood weights (Strimmer and Rambaut, 2002), and the approximately unbiased test (Shimodaira, 2002) with the –zb 10000 –au –zw options; a tree with *Paenungulates* as a polytomy was compared to the three alternate resolutions of this clade (**Figure 3A**). The 13,388 gene trees were used to infer a species tree with ASTRAL-III v.5.6.3 (Zhang et al., 2018); support for the ASTRAL species tree was also assessed with the gene-tree bootstrap approach (Simmons et al., 2019), which generated 100 pseudoreplicate datasets of the 13,388 gene trees followed by ASTRAL species tree inference on each of the 100 datasets using the *msc_tree_resampling.pl* script (Simmons et al., 2019). Gene and site concordance factors (Minh et al., 2020) were inferred with IQTREE2 v.2.2.2 COVID-edition, using the updated maximum likelihood-based method (–– scfl option) for site concordance factors (Mo et al., 2022), and the ASTRAL species tree.

**Figure 3.**
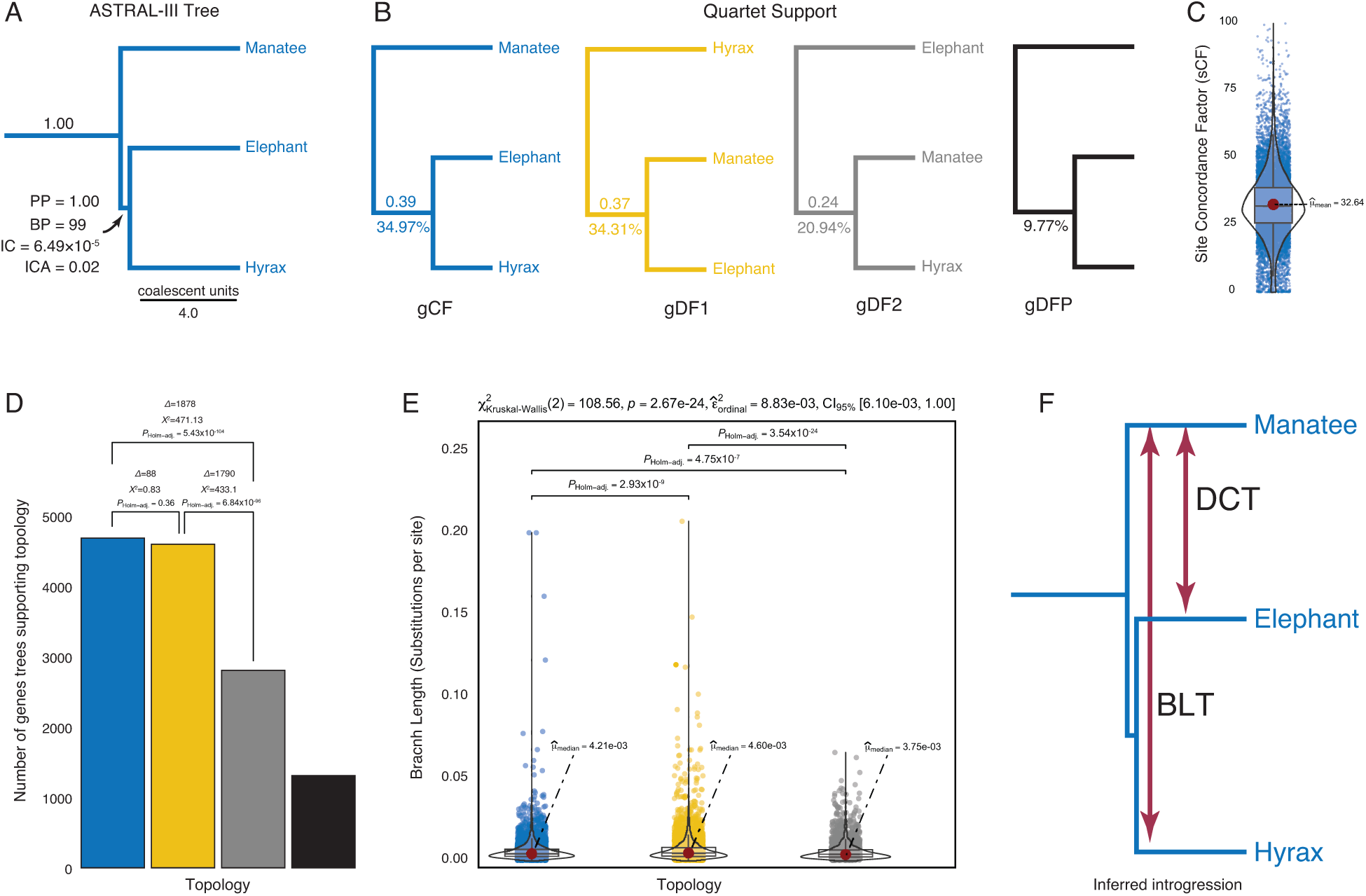
Strong support for the Paratethytheria species tree, but significant gene tree discordance and introgression. **A.** ASTRAL-III species tree, with posterior probability (PP), bootstrap proportion (BP), internode certainty (IC), and IC all (ICA) support values for the *Paratethytheria* split. Internal branch lengths in the ASTRAL species tree are shown in coalescent units (i.e., 2*N* generations), terminal branch lengths are arbitrary. **B.** Gene concordance and discordance. Gene concordance factor (gCF) for the *Paratethyeria* split, and gene discordance factors (gDF1 and gDF2) for the *Tethytheria*, and *Peritethytheria* splits are shown below each branch and quartet support for each split above. gDFP, gene discordance factor for a tree with a polyphyletic *Paenungulata*. **C.** Distribution of site concordance factors among the 13,388 genes inferred using the ASTRAL species tree. **D.** Number of gene trees supporting the *Paratethytheria*, *Tethytheria*, and *Peritethytheria* splits. The differences in the number of gene trees supporting each split (Δ), χ^2^ values, and Holm-adjusted χ^2^ *P*-values for each discordant topology test (DCT) are shown each bar. **E.** The distribution of branch lengths in gene trees supporting the *Paratethytheria*, *Tethytheria*, and *Peritethytheria* splits. Statistical significance of the difference in median branch lengths (substitutions per site) from a Kruskal-Wallis one-way ANOVA is shown above each comparison as Holm-adjusted *P*-values for the branch length test (BLT). Descriptive statistics are shown above the plot. **F.** The DCT indicates significant introgression between manatee and elephant lineages, while the BLT indicates significant introgression between the manatee and hyrax lineages.

### Characterizing incomplete lineage sorting and introgression

We used two methods to detect patterns of introgression between *Paenungualte* lineages based on the distributions of gene tree topologies and branch lengths for triplets of lineages. If the species tree is ((A, B), C), or the *Paratethytheria* resolution inferred with the ASTRAL species tree, these tests can detect introgression between A (*Probosceidea*) and C (*Sirenia*), and between B (*Hyracoidea*) and C (*Sirenia*). The discordant-count test (DCT) compares the number of genes supporting each of the two possible discordant gene trees ((A, C), B) and (A, (B, C)); in the absence of ancestral population structure, gene genealogies from loci experiencing ILS will show either topology with equal probability and ILS alone is not expected to bias the count towards one of the topologies (Huson et al., 2005; Suvorov et al., 2022). Introgression, however, will lead to a statistically significant difference in the number of gene trees, which can be evaluated with a *Χ*^2^– test (Lanfear, 2018; Suvorov et al., 2022); if there is introgression between A and C, there will be an excess of gene trees with the ((A, C), B) topology (Suvorov et al., 2022). The branch-length test (BLT) examines branch lengths to estimate the age of the most recent coalescence event (measured in substitutions per site); introgression leads to more recent coalescences than expected under the species tree topology with complete lineage sorting, while ILS shows older coalescence events (Green et al., 2010; Suvorov et al., 2022).

ILS alone does not result in different coalescence times between the two discordant topologies, and this forms the null hypothesis for the BLT that the distribution of branch lengths of gene trees supporting the ((A, C), B) and (A, (B, C)) topologies should be similar (Suvorov et al., 2022). In the presence of introgression, these branch length distributions will be skewed such that ((A, C), B) < (A, (B, C)) suggests introgression consistent with discordant topology ((A, C), B) and ((A, C), B) > (A, (B, C)) suggests introgression consistent with discordant topology (A, (B, C)). We implemented two versions of this test, one that does not scale branch lengths by total tree length and one that does (Suvorov et al., 2022); for the former, we tested the statistical significance of differences between the distribution of branch lengths with a Kruskal-Wallis one-way ANOVA while for the latter used a Mann-Whitney U test (Suvorov et al., 2022). *P*-values were corrected for multiple testing within the DCT and BLT with Holm’s method (Holm, 1979). For the scaled BLT and DCT with all trios within *Paeunungulates,* we used the *blt_dct_test.r* script (Suvorov et al., 2022).

## Results and Discussion

### Gene trees

We assembled a dataset of 13,491 orthologous protein-coding gene alignments from the genomes of 261 *Eutherian* mammals (**Supplementary Table 2**), for which we used IQTREE2 to infer the best-fitting nucleotide substtition model and maximum likelihood tree; we thus inferred 13,388 gene trees (103 tree searches failed). We found that 80% of ML gene (nucleotide) trees inferred that African elephant (*Loxodonta africana*), hyraxes (*Procavia* and *Heterohyrax*), and manatee (*Trichechus manatus*) formed a monophyletic clade (*Paenungulata*), similar to nearly all other phylogenetic analyses of *Eutherian* mammals. Relationships within *Paenungulata*, however, were almost evenly divided between the three alternative tree topologies, with the *Paratethytheria* split receiving the most support (**Figure 2A**). Branch supports assessed with the SH-like aLRT test (Anisimova et al., 2006; Guindon et al., 2010) indicate that only 252 of 13,388 genes (1.9%) had SH-like aLRT support values ≥95%, which corresponds to an 5% significance level (Anisimova et al., 2011), and these genes favor the *Tethytheria* resolution (**Figure 2A**). As expected, Pearson’s correlation tests revealed that SH-like aLRT support and branch length (substitutions per site) were positively correlated, such that longer branches tended to have higher SH-like aLRT support values (**Figure 2B**). Thus, while *Paratethytheria* received the greatest support among all genes, genes with significant phylogenetic signal (SH-like aLRT≥95%) support the *Tethytheria* resolution.

### Multispecies coalescent tree, gene and site concordance

Next, we used the 13,388 gene trees to infer a species tree with ASTRAL-III, which is statistically consistent under multispecies coalescent model (MSC) and is useful for inferring a species tree when there is significant incomplete lineage sorting. While the ASTRAL species tree supported the *Paratethytheria* split with a posterior probability of 1.00 and a gene-tree bootstrap support of 98.7% (**Figure 3A**), there can be significant underlying gene tree discordance even for branches with near 100% support (Jarvis et al., 2014; Pease et al., 2016; Salichos and Rokas, 2013; Vanderpool et al., 2020). For example, the internode certainty (IC), which corresponds to the magnitude of conflict between the two most likely splits among gene trees (Salichos et al., 2014) for the *Paratethytheria* split is 6.49×10^-5^ (**Figure 3A**); IC values near 1 indicate the absence of conflict for the split, whereas IC values near 0 indicate equal support for each split and hence maximum conflict (Salichos et al., 2014). Note that the maximum IC, corresponding to one more tree supporting *Paratethytheria* over *Tethytheria* is 3.35×10^-8^. Similarly, the internode certainty all (ICA), which corresponds to the magnitude of conflict between all splits among gene trees (Salichos et al., 2014) is 0.02 (**Figure 3A**); ICA values at or near 1 indicate the absence of any conflict among splits, whereas ICA values near 0 indicate that one or more conflicting splits have almost equal support (Salichos et al., 2014). Thus, while there is strong support for the *Paratethytheria* resolution in the MSC species tree, there is also significant gene tree discordance for this split.

We next used IQTREE2 (Nguyen et al., 2015) to infer gene (gCF) and site concordance (sCF) factors for each branch in the ASTRAL species tree; gCF and sCF are the fraction of genes and sites that are in agreement with the species tree for any particular branch (Minh et al., 2020; Mo et al., 2022). Consistent with significant phylogenetic uncertainty among gene trees, the gCF was only 34.97% for the *Paratethytheria* split, whereas the discordance factor (the proportion of trees supporting alternative resolutions) was 34.31% for the *Tethytheria* split and 20.94% for the *Peritethytheria* split; 9.77% of gene trees were discordant due to polyphyly (**Figure 3B**). Similarly, quartet support for the *Paratethytheria* was 0.39, 0.37 for the *Tethytheria* split, and 0.24 for the *Peritethytheria* split (**Figure 3B**). Site concordance factors inferred for each gene using the ASTRAL species tree were also very low, with a median of 32% and a mean of 33%, indicating a high rate of discordance among sites (**Figure 3C**). Thus while the species tree strongly supports *Paratethytheria*, concordance factors suggest that most genes and sites support a different relationship. Indeed, there is almost perfect discordance between the *Paratethytheria* and *Tethytheria* splits.

### Discordance from incomplete lineage sorting and introgression

If the discordance among gene trees results from incomplete lineage sorting (ILS), the number of gene trees supporting the two discordant topologies should be roughly equal (Huson et al., 2005). In contrast, introgression, or a combination of ILS and introgression, will cause one of the two discordant topologies to be more frequent (Suvorov et al., 2022); the difference between the number of gene trees supporting alternative resolutions can be compared with a *Χ*^2^–test (Lanfear, 2018; Suvorov et al., 2022), and is called the discordant-count test (DCT)^3^. We found that the number of gene trees supporting *Paratethytheria* and *Tethytheria* was significantly greater than the number supporting *Peritethytheria*, but there was not a significant difference between *Paratethytheria* and *Tethytheria* (**Figure 3D**). These data indicate an excess of gene trees supporting *Tethytheria*, consistent with introgression from *Sirenia* into *Proboscidea* (**Figure 3F**).

ILS alone will not result in different coalescence times between the two discordant trees, while introgression will result in more recent coalescences than expected under the species tree topology with complete lineage sorting (Suvorov et al., 2022); the branch length test (BLT) compares the distribution of branch lengths for each gene tree with discordant topologies, with the expectation that they are similar under ILS alone (Suvorov et al., 2022). The distribution of branch lengths between gene trees supporting *Paratethytheria* and *Tethytheria* was significantly different than *Peritethytheria*, as was the difference between *Paratethytheria* and *Tethytheria* (**Figure 3E**). We also used the scaled BLT of Suvorov et al. (2022), as well as their approach of testing trios of lineages for the DCT, rather than our approach of testing elephant, manatee, and the *Procavia*-*Heterohyrax* stem-lineage, and found similar results (**Supplementary Table 3**). These data are consistent with introgression from *Sirenia* into *Hyracoidea* (**Figure 3F**)

### Other sources of discordance

Technical noise including base composition variation, gene length, low phylogenetic signal, and model misspecification, among many others, also contributes to gene tree discordance. Longer genes tend to have more phylogenetically informative sites than shorter genes, but while there were statistically significant differences in the length of genes that support each of the three splits the effect sizes were small (**Supplementary** Figure 1A). Genes with high GC content can have poor phylogenetic resolution because they can have many homoplasies at hypermutable CpG sites and more frequent biased gene conversion (Romiguier et al., 2013). However, GC-content was similar between genes supporting the three splits (**Supplementary** Figure 1A). Indeed, a tree inferred from the third of genes with the lowest GC content (n=4,444) concatenated into a supermatrix, with separate a GTR6 model for each gene partition, supported the *Paratethyeria* split (**Supplementary** Figure 2A), as did consensus trees when genes were binned into quartiles based on GC content (**Supplementary** Figure 2B). These data indicate that our results are robust to noise from GC content variation across genes. Genes supporting the *Paratethytheria* split had statistically higher SH-like aLRT scores, sCFs, and longer branch lengths than genes supporting the other splits, again, however, the effect sizes were small (**Supplementary** Figure 1A). There also were no meaningful correlations between sCF (calculated from the ASTRAL species tree with the *Paratethytheria* split) and gene length, GC-content, SH-like aLRT score, or branch length, or between SH-like aLRT score and gene length, GC-content, or branch length (**Supplementary** Figure 1B). Thus, while each of these factors may contribute to gene tree discordance, their effects are likely small (but they may also be additive).

### Tree topology tests

Few studies have explicitly tested whether gene trees support alternative resolutions of *Paenungulates* (Liu et al., 2023). Therefore, we used IQTREE2 to perform several tree topology tests, including the Kishino-Hasegawa test (Kishino and Hasegawa, 1989), the Shimodaira-Hasegawa tests (Shimodaira and Hasegawa, 1999), the weighted Kishino-Hasegawa and Shimodaira-Hasegawa tests, the approximately unbiased (AU) test (Shimodaira, 2002), the expected likelihood weights (Strimmer and Rambaut, 2002), and the bootstrap proportion (Kishino et al., 1990). A tree with *Paenungulates* as a polytomy was compared to the three alternate resolutions of this clade (**Figure 4A**); therefore, these methods test if there is an improvement in the likelihood score of the three alternative topologies compared to the polytomy tree (Shen et al., 2017). We found that the likelihood differences (deltaL) between the polytomy and three alternative trees were generally very small for most genes (**Figure 4B**) and that the suite of tree topology tests agreed the worst tree was *Peritethytheria* and the best tree was the *Paratethytheria* (**Figure 4C**). However, only 8-10% of genes statistically rejected the polytomy tree (**Figure 4C**). Thus, very few genes have statistically significant phylogenetic signal to reject the *Paenungulate* polytomy tree.

**Figure 4.**
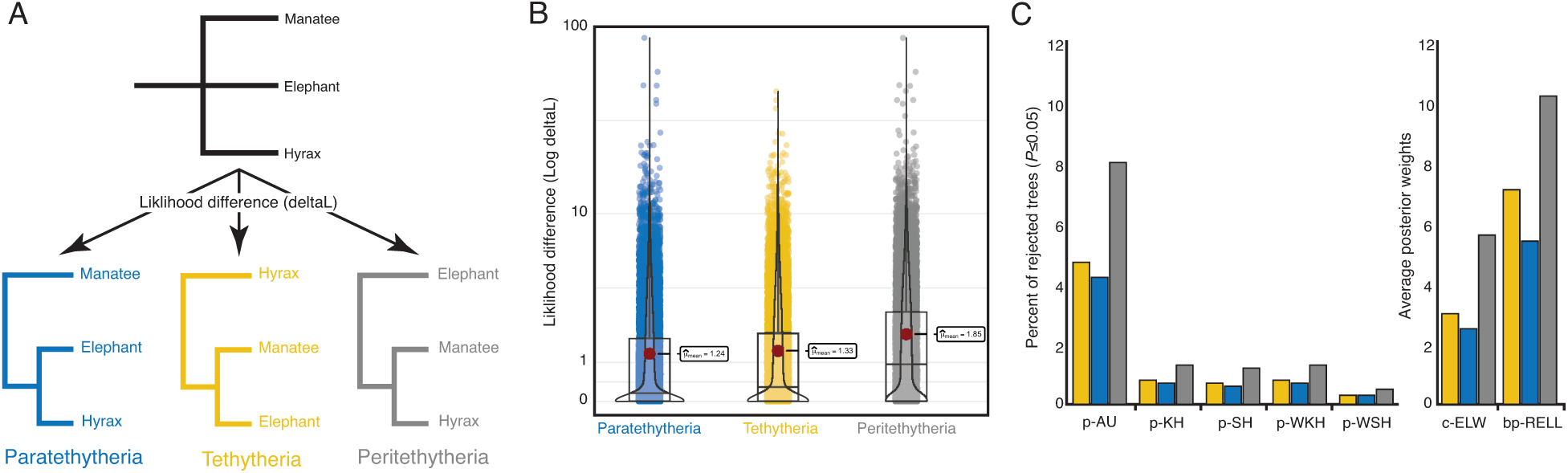
Tree topology tests do not support resolution of the *Paenunulate* polytomy. **A.** The reference tree used for topology tests was the ASTRAL species tree with *Paenungulates* collapsed to polytomy, the likelihood of which was compared to the likelihood of the ML tree inferred for each gene. **B.** Likelihood difference between the *Paenungulate* polytomy tree and ML gene tree for each gene. The likelihood difference is shown for each gene as jittered dots, overlaid violinplots, and boxplots show the data distribution. **C.** Percent of genes that reject each alternative tree, i.e., did not have a statistically significant improvement in likelihood score compared to the polytomy tree (*P*≤0.05), inferred from the Kishino-Hasegawa (p-KH), the Shimodaira-Hasegawa (p-SH), the weighted Kishino-Hasegawa (p-WKH), the weighted Shimodaira-Hasegawa (p-WSH), and the approximately unbiased test (p-AU) tests, and the posterior weights from the expected likelihood weight (c-ELW) from the bootstrap proportion (bp-RELL) tests.

### Caveats and limitations

Inferring reliable gene and species trees from phylogenomic data, and thus sorting among competing scenarios of gene tree inference error, incomplete lineage sorting, and introgression (Hibbins and Hahn, 2021) is fraught with *ad hoc* decisions, methodological limitations, and technical challenges. For example, we have focused on inferring phylogenetic relationships from protein-coding genes, for which we could generate high-quality alignments for 261 *Eutherian* mammals. While our dataset is extensive, including 13,388 genes with a total alignment length of 19,968,803 bp, the coding fraction of the genome is relatively small and may have less or different phylogenetic signal than non-coding regions (Foley et al., 2015, 2023; Literman and Schwartz, 2021; William J Murphy et al., 2001). Despite this potential limitation, our observation that a slight majority of gene trees (n=88) support the *Paratethytheria* split is similar to a recent study based on 411 kb of genome-wide nearly neutral sites and non-coding sites (Foley et al., 2023); however, our results differ in supporting *Tethetheria* rather than *Peritehytheria* as the second most likely split. These data suggest that systematic differences in phylogenetic information content between coding and non-coding (or genic and intra/inter-genic) regions are unlikely to bias our results, but it remains a possibility.

Other technical sources of error, such as misalignment, model misspecification, and long branch attraction (LBA), could also bias our gene tree inferences and downstream analyses. For example, alignment error leads to statistical inconsistency of phylogenetic methods, manifesting in nucleotide alignments as gross substitution model misspecification and LBA artifacts (Hossain et al., 2015; Ogden and Rosenberg, 2006). We attempted to mitigate the effects of alignment error through stringent alignment criteria; however, assessing the impact of such error (Landan and Graur, 2008; Sela et al., 2015) in phylogenomic scale datasets is computationally impractical. Similarly, we inferred gene trees with the best substitution model for each gene to minimize the effects of model misspecification. Still, computational limitations prevented us from using more complex models that account for model variation within genes, such as mixture models (Lartillot and Philippe, 2004; Le et al., 2012; Quang et al., 2008; Wang et al., 2008) and site-specific frequency models (Quang et al., 2008; Wang et al., 2018), and processes like heterotachy, i.e., rate variation across sites and lineages (Crotty et al., 2019; Kolaczkowski and Thornton, 2008, 2004), which may be common in coding regions. However, whether model selection is an essential step in phylogenetic inference is debated (Abadi et al., 2019; Doko and Liu, 2023; Gerth, 2019; Wu et al., 2014), and likelihood methods may be robust to all but the most extreme model violation (Naser-Khdour et al., 2021). Indeed, the wrong model may infer the correct tree more often than the true model (Bruno and Halpern, 1999; Yang, 1997). Thus, we do not believe that model misspecification negatively impact our results. Still, we cannot exclude the possibility that extreme convergence following the very rapid diversification of *Paenungulates* has led to long-branch attraction artifacts or that heterotachy has affected gene tree inference for some genes. For example, in a previous study on another rapid radiation event, the branching order of the Hox cluster duplications, we found that genes with internal branch lengths < 0.02 were significantly impacted by LBA artifacts (Lynch and Wagner, 2009); only 5.4% of genes had *Paratethytheria*, *Tethetheria*, and *Peritehytheria* stem-lineages longer than 0.02, and the majority of these (50.07%) support the *Tethytheria* split, thus LBA artifiacts are a possible source of gene tree error. Given that the *Paratethyeria* resolution is supported by only 88 genes more than the *Tethytheria* resolution, incorrect tree inference in even a few genes could adversely affect our results.

## Conclusions

Resolving the phylogenetic relationships within *Paenungulates* has been remarkably challenging, with different studies and data supporting different resolutions. Our data indicate that 92.7% protein-coding genes lack significant phylogenetic signal to reject a *Paenungulate* polytomy and that the branching order of *Paenungulates* is obscured by large-scale gene tree error, incomplete lineage sorting, and introgression. This combination of factors suggests that previous molecular phylogenetic studies, even relatively large multigene studies, were biased by stochastic gene sampling, contributing to the many conflicting resolutions of relationships within *Paenungulates*. In contrast, our study includes 13,388 protein-coding genes, representing ∼67% of the coding genome and nearly 20 million base pairs, and is unlikely to be biased by gene sampling artifacts. Thus, we conclude that the combination of systematic biases in gene tree estimation caused by very rapid divergence leading to low phylogenetic information content and significant gene tree uncertainty, high levels of ILS, and introgression combine to create a perfect storm scenario (Cai et al., 2020) for inferring the branching order within *Paenungulates*. Therefore, this clade is likely as close to a real, or at least likely unresolvable, polytomy as possible.

## Acknowledgments

VJL is partially supported by NIH award 1R56AG071860-01. Computational support was provided by the Center for Computational Research at the University at Buffalo.

## Data availability

Alignments (https://doi.org/10.5061/dryad.5dv41nsbc) as fasta files and IQTREE result output files for each gene (https://doi.org/10.5061/dryad.j0zpc86n3) are available from Dryad.

**Supplementary Figure 1.**
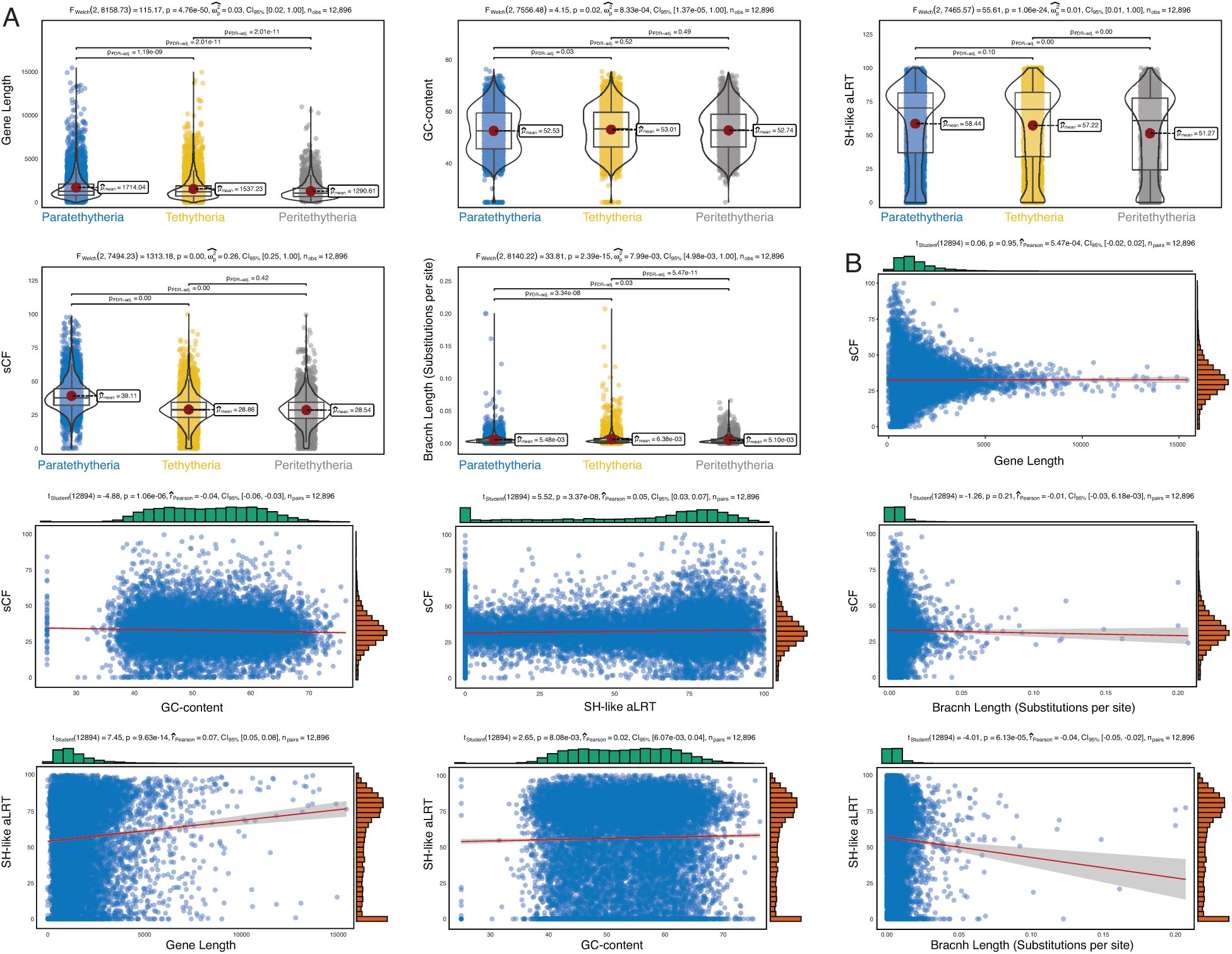
Other sources of phylogenetic discordance among Paenungulates. **A.** Stripchart/violin/boxplots showing differences in mean gene length, GC-content, SH-like aLRT score, sCF, and branch length between genes with gene trees that support the *Paratethytheria*, *Tethytheria*, and *Pertethytheria* splits. Summary statistics and FDR-adjusted *P*-values from pair-wise Games-Howell tests are shown. **B.** Scatterplots showing the correlation between variables in panel A; sCF, SH-like aLRT, and branch length were calculated from the ASTRAL species with the *Paratethytheria* split. Summary statistics from Pearson’s correlation coefficients are shown above reach split, and side histograms show the distribution of values for each gene.

**Supplementary Figure 2.**
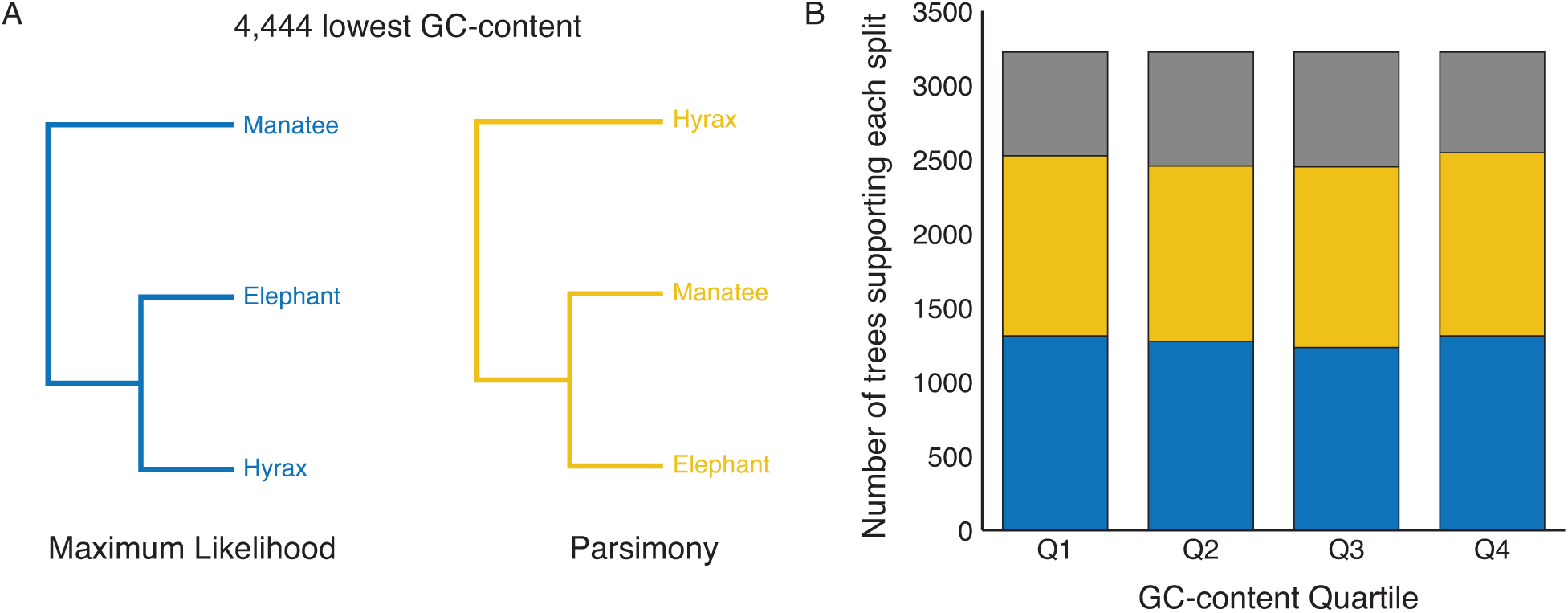
GC content variation across genes is an unlikely source of gene tree discordance. **A.** Maximum likelihood and parsimony trees inferred from a concatenated supermatrix of the third of genes with the lowest GC content (n=4,444). A GTR substitution model was used for each gene partition in the supermatrix. **B.** Number of gene trees supporting each split when genes are binned by GC content quartiles. The *Paratethytheria* split is supported by the greatest number of gene trees in each quartile.

---

**Supplementary Table 1.**
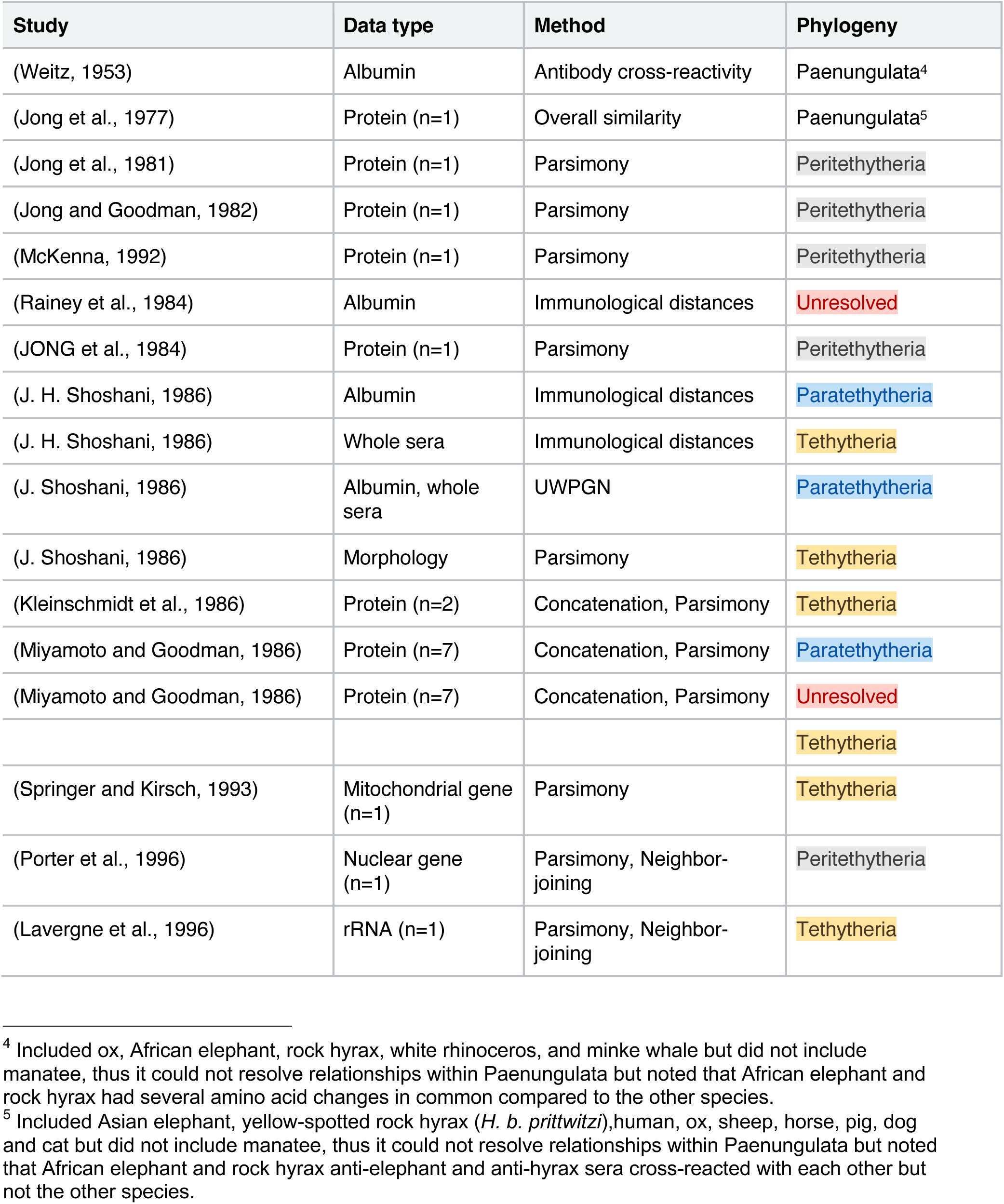
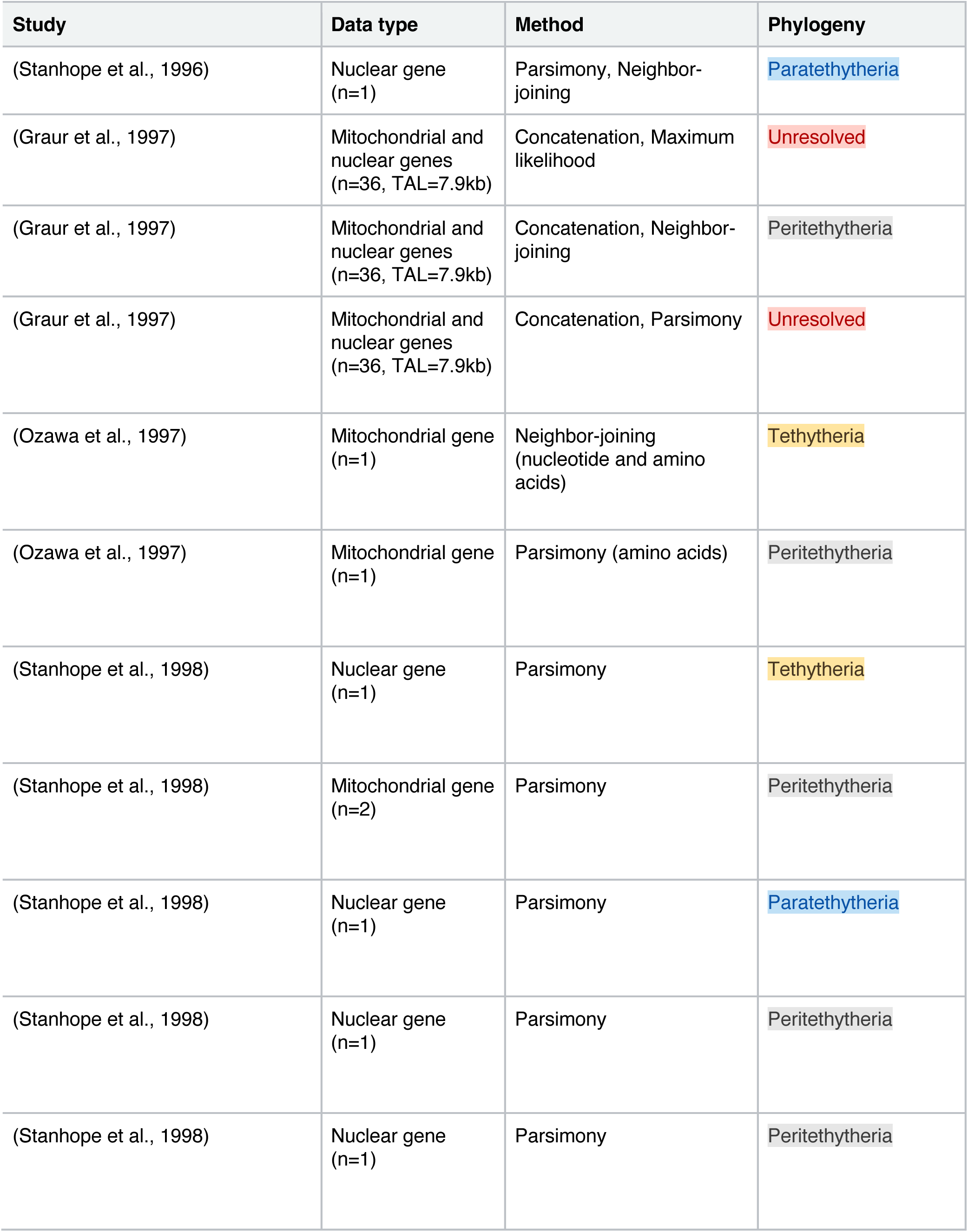
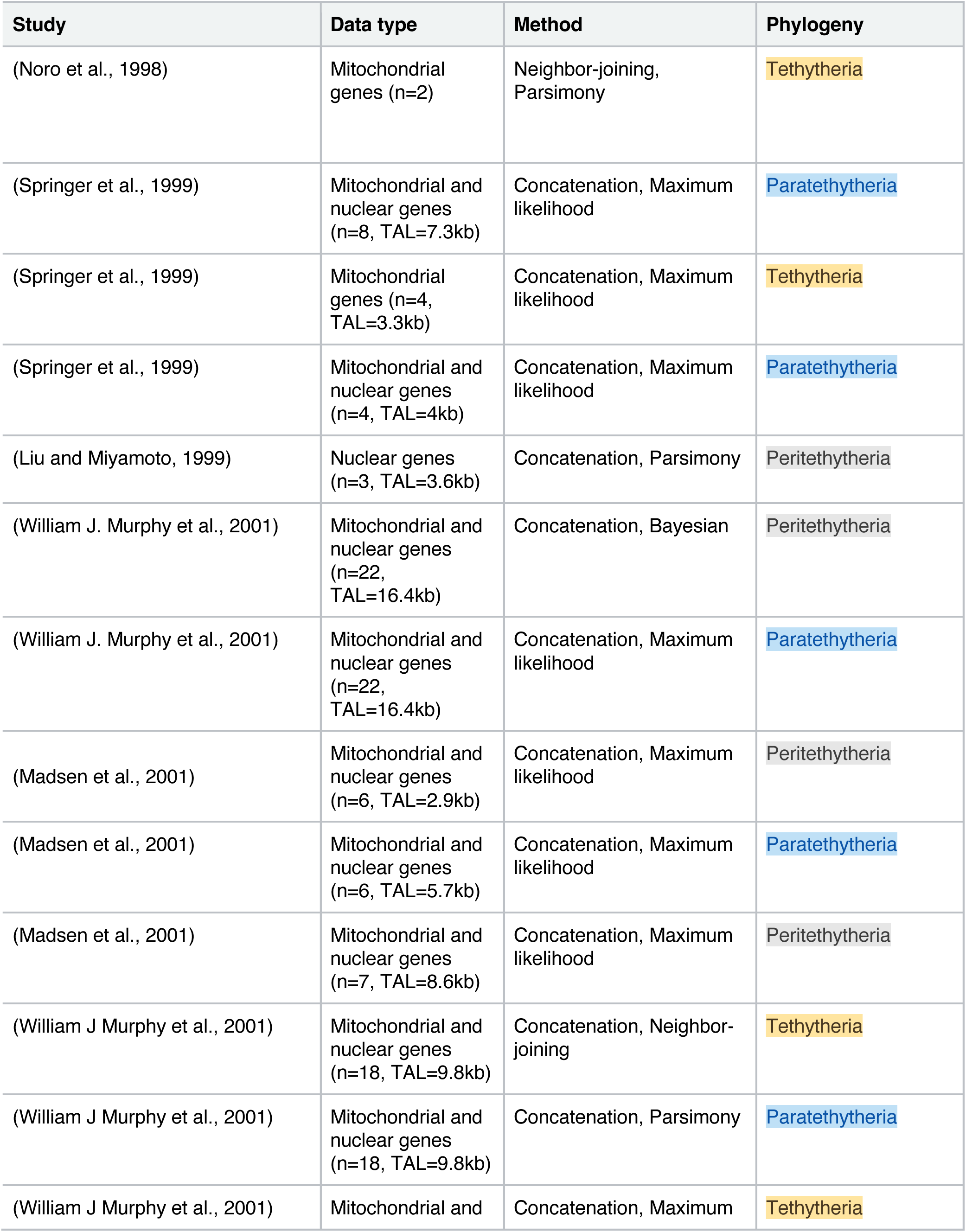
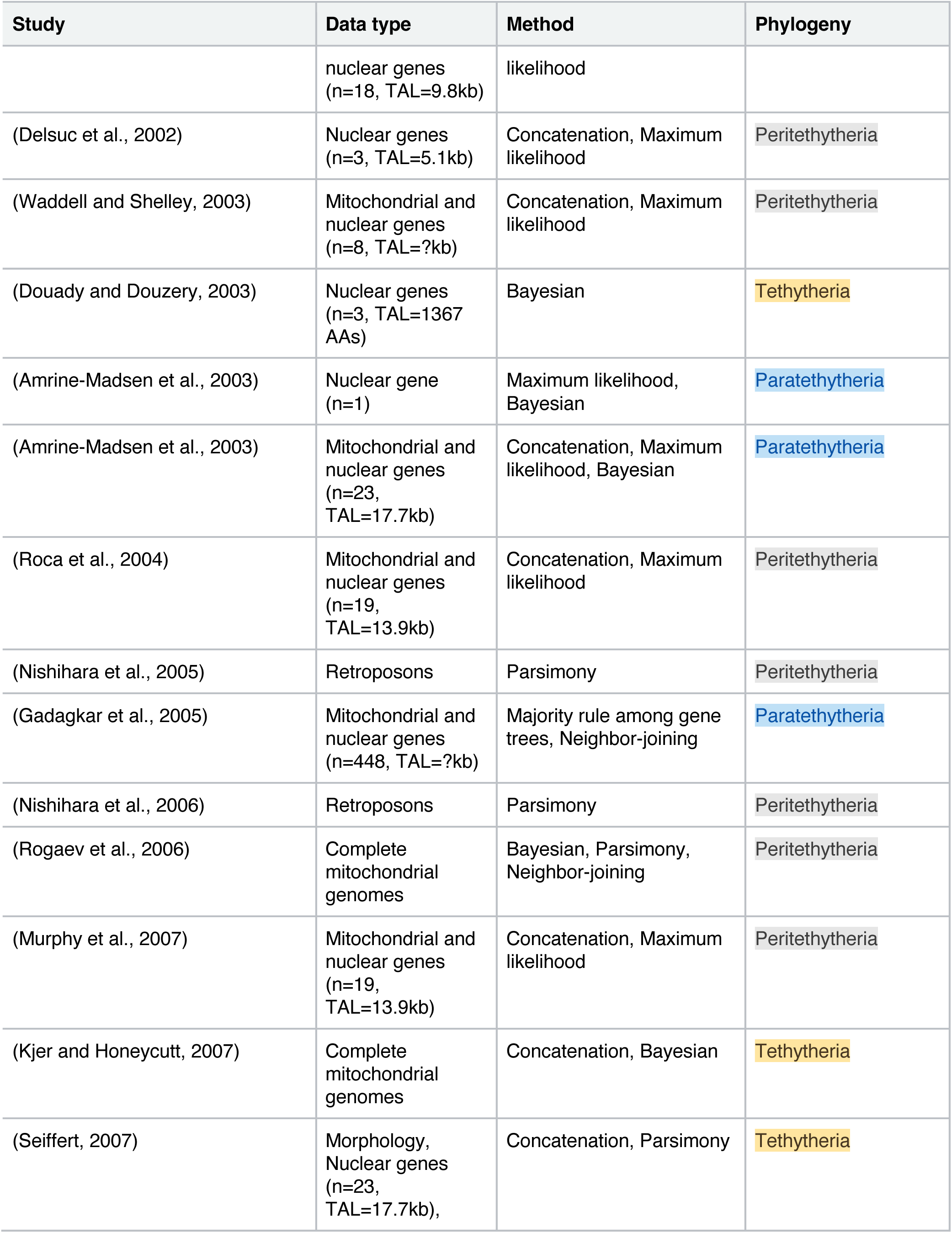
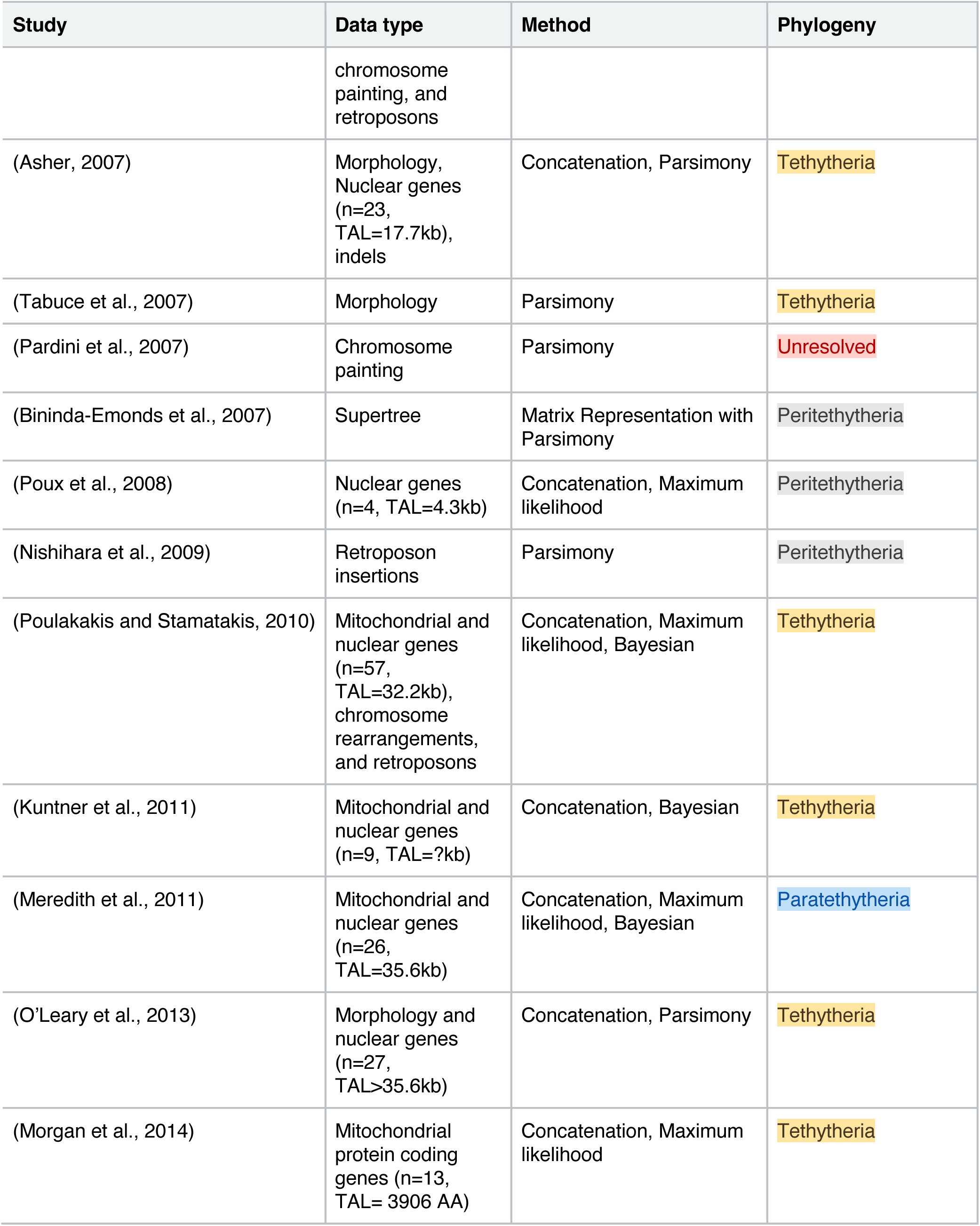
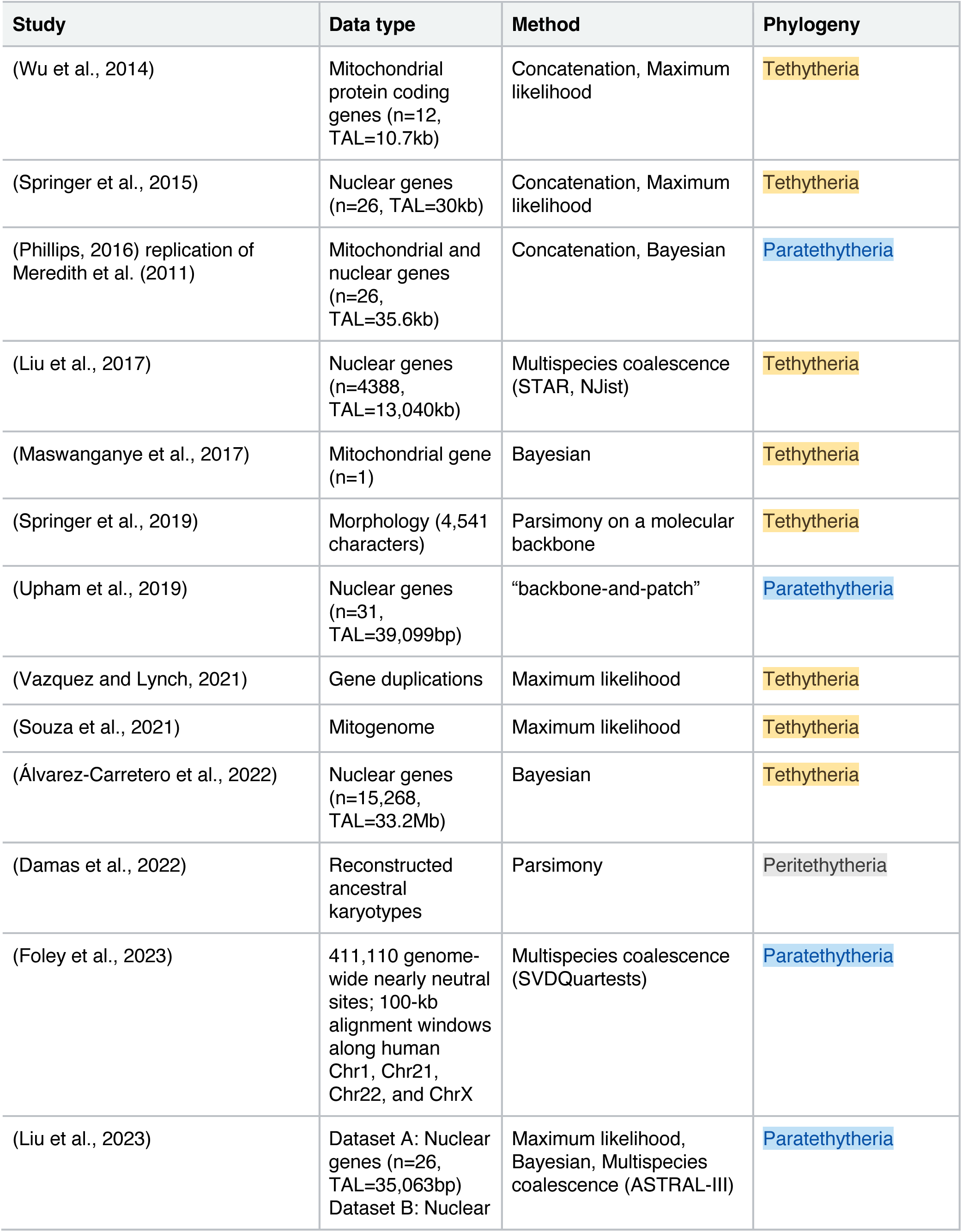
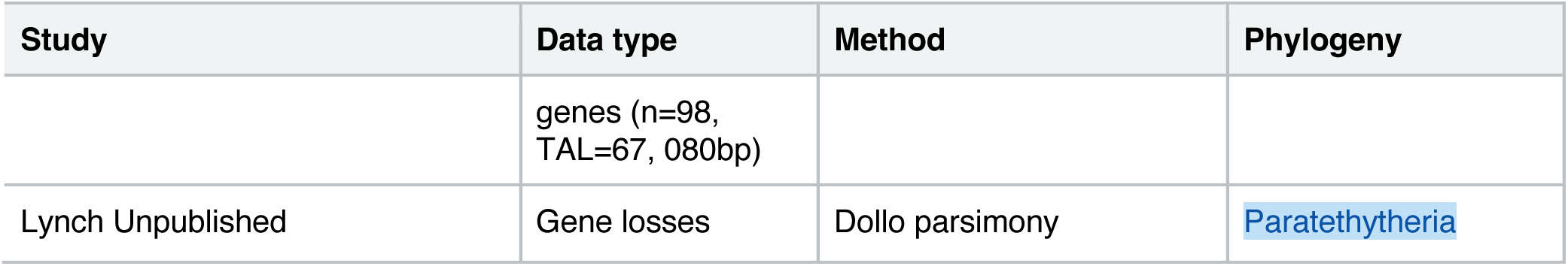
Molecular phylogenetic studies that have included *Paenungulata*. If the study has more than 10 genes, total alignment length (TAL) is shown.

1 We follow the formatting suggestion of (Thines et al., 2020) and placed scientific names at all taxonomic ranks in italics to facilitate their easy recognition.

2 To our knowledge, no in/formal names have been proposed for the alternative *Proboscidea*+*Hyracoidea* or *Sirenia*+*Hyracoidea* resolutions of *Paenungulata.* Therefore, we refer to the *Proboscidea*+*Hyracoidea* clade as *Paratethytheria*, after the Paratethys sea, which was repeatedly disconnected from and reconnected with the Tethys sea, and the *Sirenia*+*Hyracoidea* clade as *Peritethytheria*, to also reflect the origin of these lineage around the Tethys sea.

3 As initially proposed (Huson et al., 2005), the significance of the difference in number between the two discordant trees can be compared with the Azuma-Hoeffding inequality (Alon and Spencer, 2023). Using this method, the number of gene trees supporting *Paratethytheria* (Δ=1878, *P_Holm-adj._*=4.88×10^-26^) and *Tethytheria* (Δ=1790, *P_Holm-adj._*=4.05×10^-23^) was significantly greater than the number supporting *Peritethytheria*, but there was not a significant difference between *Paratethytheria* and *Tethytheria* (Δ=88, *P_Holm-adj._*=0.20).

